# Multi-Trait Machine and Deep Learning Models for Genomic Selection using Spectral Information in a Wheat Breeding Program

**DOI:** 10.1101/2021.04.12.439532

**Authors:** Karansher S. Sandhu, Shruti S. Patil, Michael O. Pumphrey, Arron H. Carter

**Author notes:** **Correspondence:** Dr. Arron H. Carter.

## Abstract

Prediction of breeding values and phenotypes is central to plant breeding and has been revolutionized by the adoption of genomic selection (GS). Use of machine and deep learning algorithms applied to complex traits in plants can improve prediction accuracies in the context of GS. Spectral reflectance indices further provide information about various physiological parameters previously undetectable in plants. This research explores the potential of multi-trait (MT) machine and deep learning models for predicting grain yield and grain protein content in wheat using spectral information in GS models. This study compares the performance of four machine and deep learning-based uni-trait (UT) and MT models with traditional GBLUP and Bayesian models. The dataset consisted of 650 recombinant inbred lines from a spring wheat breeding program, grown for three years (2014-2016), and spectral data were collected at heading and grain filling stages. MT-GS models performed 0-28.5% and −0.04-15% superior to the UT-GS models for predicting grain yield and grain protein content. Random forest and multilayer perceptron were the best performing machine and deep learning models to predict both traits. These two models performed similarly under UT and MT-GS models. Four explored Bayesian models gave similar accuracies, which were less than machine and deep learning-based models, and required increased computational time. Green normalized difference vegetation index best predicted grain protein content in seven out of the nine MT-GS models. Overall, this study concluded that machine and deep learning-based MT-GS models increased prediction accuracy and should be employed in large-scale breeding programs.

**Core Ideas:**

1. Potential for combining high throughput phenotyping, machine and deep learning in breeding.
2. Multi-trait models exploit information from secondary correlated traits efficiently.
3. Spectral information improves genomic selection models.
4. Deep learning can aid plant breeders owing to increased data generated in breeding programs

## Introduction

Quantitative genetics theory was proposed by Sir Ronald Fisher a century ago and established the infinitesimal model (Fisher, 1918). This theory was developed without the direct use of genotypic data and persisted for decades. With the advent of sufficient genome-wide markers paired with an infinitesimal model of quantitative genetics theory, Meuwissen et al. (2001) were the first to propose the term genomic selection (GS) for predicting breeding values in animals and plants. GS uses estimates of marker effects based on model development from a related population genotyped and phenotyped for the trait of interest. The genome-wide marker effects are then used for predicting the genomic estimated breeding values (GEBVs) of a new population, which is only genotyped (Heffner et al., 2010). GS has been extensively researched in plant breeding, focusing on optimizing marker density, training population size, family relatedness, heritability of the trait, and GS model utilized (Shengqiang et al., 2009). Most of the GS models evaluated in plant breeding have been uni-trait (UT), where single traits are predicted (Lozada & Carter, 2019; Sun et al., 2019). Recently, breeders have moved to adopt multi-trait (MT) GS models because predictions are required for multiple traits simultaneously and their combination may improve prediction accuracies (Jia & Jannink, 2012).

MT-GS models leverage shared genetic information between correlated traits (Jia & Jannink, 2012) to predict various traits simultaneously by utilizing the same set of predictors, assuming the presence of some structure in the captured output (Bhatta et al., 2020). Improved prediction accuracy of MT-GS models over UT-GS is attributed to the correlation among the training population and between the traits. Furthermore, MT-GS models are more interpretable and require less computational time than a series of UT-GS models (Montesinos-López et al., 2018a). MT models are intensively employed in other fields such as data mining, forest management, energy forecasting, and ecological modelling (Voyant et al., 2017). Jia & Jannink (2012) showed that prediction accuracy improved for primary traits with low heritability in barley (*Hordeum vulgare* L.) when a secondary correlated trait is used in MT-GS models. Sun et al. (2019) demonstrated the increase in GS prediction accuracy for wheat (*Triticum aestivum* L.) grain yield when secondary traits such as normalized difference vegetation index (NDVI) and canopy temperature were included in MT-GS models. Bhatta et al. (2020) compared UT- and MT-GS models for predicting end-use quality traits in barley and concluded that MT-GS models have better performance under both within and across environment predictions. Mixed model approaches utilizing Bayesian and genomic best linear unbiased predictor (GBLUP) are most commonly used in plant breeding programs (Endelman, 2011; Pérez & Campos, 2014).

GBLUP is a frequently used MT-GS models in plant breeding, which uses marker-based relationship matrix for predicting the performances of genotypes (Endelman, 2011; VanRaden, 2008). Several Bayesian MT-GS models (Bayes A, Bayes Lasso, Bayes B, and Bayes Cpi) are also available, which assume a prior distribution during the training process, and hence separate models are required to optimize different traits (Pérez & Campos, 2014). Bayes Lasso follows the double exponential prior distribution for performing the continuous shrinkage and variable selection (Tishbirani, 1996). Additionally, Bayes Lasso applies a long tail student t distribution to the marker effects. Bayes A, Bayes B, and Bayes Cpi use the scaled-t, gaussian mixture (point mass at zero with gaussian distribution), and scaled-t mixture (point mass at zero with scaled-t distribution) prior distribution, respectively, during the model training (Pérez & Campos, 2014). These models assume that all markers do not contribute to the total genetic variance. Bayesian models are also computationally intensive due to Monte Carlo Markov Chain utilization for estimating the marker effects. Bayesian models are known as parametric due to the assumption of the prior relationship among features and predictors, which do not model gene by gene and higher-order interactions during the estimation of marker effects (Gianola et al., 2006; Montesinos-López et al., 2019). Hence, recently developed machine and deep learning tools provide an opportunity for the selection of the GS model.

The increasing adoption of high throughput genotyping and phenotyping tools by plant breeders has increased data generation tremendously, which requires the adoption of analytical methods used in other disciplines for complex datasets. Machine and deep learning models have been explored in previous studies for prediction in UT-GS models and have demonstrated mixed results (Bellot et al., 2018; Ma et al., 2017). Random forest (RF) and support vector machine (SVM) are commonly used ensemble machine learning models for the GS context. RF is an ensemble machine learning tool used for predicting the output by averaging the results of an extensive collection of identically distributed decision trees applied to bootstrapped samples of the training data. RF is better than the other tree-based methods like decision tree regression and bagging, as it tries to reduce the correlation between the subsets of tree by averaging results from those trees for final predictions. The reduction of correlation among the independent trees and averaging performance of the trees aids in reducing overfitting and increasing the prediction accuracy. The important hyperparameters for RF model training include the number of trees, the number of features sampled for each iteration, the importance of each feature, and the depth of the trees (Hastie et al., 2009). SVM provides flexibility for fitting the model as it fits the best regression line by allowing a specific acceptable error in the model. Optimization of SVM models requires finding a regression line that deviates from the real line by no greater than a value called maximum error (*ϵ*), and at the same time the line should be flat as possible (Smola & Scholkopf, 2004).

Deep learning is the branch of machine learning that uses an artificial neural network as a prediction tool, and needs to be explored in GS owing to the plethora of data accumulated in breeding programs (Lecun et al., 2015; Samuel, 2001). Deep learning models explore the relationship between input and output variables using a combination of neurons and hidden layers to form a network similar to the biological network of neurons in the human brain. Deep learning models use different non-linear activation functions with a large number of layers, and data is transformed along with each layer to obtain the best fit for different genetic architectures (Angermueller et al., 2016). Often used deep learning models in plant breeding are multilayer perceptron (MLP) and convolutional neural network (CNN) (Pérez-Enciso & Zingaretti, 2019). Optimization of hyperparameters is required to achieve the best deep learning model performance, which is the most computationally intensive step (McKay, 1992). The most essential hyperparameters include the type of activation function, activation rate, regularization parameter, number of epochs, number of hidden layers, dropout, and stopping criteria (Pérez-Enciso & Zingaretti, 2019). Hyperparameters can be selected by using one of the four methods, namely grid search, latin hypercube sampling, random search, and optimization (McKay, 1992).

MLP is considered a feed-forward neural network, and consists of three input, hidden, and output layers. The first layer is the input layer, which receives the DNA marker information. Each neuron of the hidden layers has its characteristic weight and transforms the previous layer’s data using various linear and non-linear activation functions (Lecun et al., 2015). The weight parameters and other hyperparameters are optimized using the training data using either backpropagation or stochastic gradient (Cho & Hegde, 2019). The number of the output layer is equal to the total number of response variables, depending upon the tested model. The output of a layer also depends upon the weighted average transformation of neurons from the previous layer with associated bias. CNN is a special type of neural network used for input features having a specific pattern, such as linkage disequilibrium among markers distributed along a linear chromosome. The hidden layer in MLP is replaced with multiple layers in CNN, such as convolutional, pooling, fully connected, and dense layers (Lecun et al., 2015). As opposed to neurons, CNN uses the kernels or filters in convolution layers for capturing the hidden information. A filter consists of predefined marker interval windows having the same weights. This filter is moved continuously across the input data for obtaining the weight for each window for computing the locally weighted sum. The pooling layer follows the first convolutional layer and is used for dimensionality reduction (Pérez-Enciso & Zingaretti, 2019). This layer merges the output of filters from the convolutional layer using either mean, minimum, or maximum to smoothen the results. Dropout and activation functions are employed after the convolutional and pooling layer (Pook et al., 2020).

High-throughput phenotyping applications include spectral reflectance values obtained from plants to provide information about various physiological processes and have been used in wheat (Babar et al., 2006), rice (*Oryza sativa* L.; Zheng et al., 2018) maize (*Zea mays* L.; Aguate et al., 2017), barley (Barmeier et al., 2017), and sorghum (*Sorghum bicolor* L.; Habyarimana et al., 2020). Different vegetation indices or spectral reflectance indices (SRI) can be extracted by measuring reflection from plants. These SRI aid in the indirect selection of primary traits (grain yield or grain protein content) in wheat due to their moderate to high genetic correlation and high heritability compared to primary traits (Crain et al., 2018). Commonly utilized indices are NDVI, photochemical reflectance index (PRI), normalized water index (NWI), and green-NDVI (GNDVI), which provide information about plant biomass, photochemical pigments, plant water stress, and nitrogen status (Gitelson et al., 1996; Peñuelas et al., 1994). These indices have been used in covariate and MT-GS models to predict grain yield in wheat and demonstrate improvement in prediction accuracy (Lozada & Carter, 2019; Sun et al., 2019). To the best of our knowledge, deep learning models have not been explored for MT-GS in wheat for predicting grain yield and grain protein content using spectral information as secondary traits.

SRI derived from wheat has been reported to correlate to grain yield, biomass, and drought tolerance in spring wheat (Gizaw et al., 2018). Grain yield and grain protein content are important selection traits in spring wheat breeding programs and are complicated by the negative correlation between them. However, GS and SRI provide an alternative for selecting these two traits simultaneously. Our previous study observed that inclusion of secondary correlated traits results in improved prediction accuracy for grain yield and grain protein content by using the rrBLUP GS model. Grain protein content and grain yield were accurately predicted when spectral data was collected at heading and grain filling stages, respectively (Sandhu et al., 2021b). Similarly, we observed that deep learning based GS models improve prediction accuracy by 3-5% in different agronomic traits in wheat (Sandhu et al., 2021a). The main objectives of this study were to 1) Optimize different MT machine and deep models for predicting grain yield and grain protein content in wheat, 2) Compare the performance of MT-GS and UT-GS models, and 3) Compare the performances of MT-GS mixed models with machine and deep models under cross-validation and independent validation scenarios.

## Materials and Methods

### Field data and plant material

The data set used in this study consisted of 650 recombinant inbred lines from a nested association mapping population of spring wheat (Blake et al., 2019). The field trial was planted for three years (2014-2016) at Spillman Agronomy Farm, Pullman, WA. For detailed information about the population, field trials, and traits evaluated see Sandhu et al. (2021a,b). In brief, field trials were planted in a modified augmented design with 15-20% of the plots assigned to three replicated check cultivars. Grain protein content (%) and grain yield (t/ha) was collected using a Perten DA 700 NIR analyzer (Perkin Elmer, Sweden) and Wintersteiger Nursery Master combine (Ried im Innkreis, Austria).

Spectral reflectance at 16 different bands between the 430 and 980 nm wavelengths was collected with a handheld CROPSCAN multi-spectral radiometer at heading (Feekes growth stages 10.1) and grain filling stages (Feekes growth stages 11.1) (Large, 1954). Data from the CROPSCAN was processed with CROPSCAN MSR software, and six different SRI were derived. These indices were NDVI, PRI, NWI, anthocyanin reflectance index (ARI), normalized chlorophyll pigment ratio index (NCPI), and GNDVI (Peñuelas et al., 1994; Prasad et al., 2007; Rouse et al., 1972). Detailed information about these SRI and the physiological traits they explain is provided in **Supplementary Table 1**.

### Genotyping

The population was genotyped using genotyping by sequencing and Illumina 90K SNP chip array (Poland & Rife, 2012). Initial genotyping data consisted of 73,345 high-quality polymorphic markers anchored to the Chinese Spring RefSeqv1 reference map (Jordan et al., 2018). Detailed procedure about genotyping, SNP calling, map construction, and filtration is described in previous publications (Sandhu et al., 2021a). Quality filtering involved removing monomorphic markers and markers missing more than 20% of the genotyping data. RILs missing phenotyping data in one environment and 10% genotyping data were discarded. Finally, a minor allele of < 0.05 was used, resulting in 635 RILs having 40,038 polymorphic markers.

### Statistical analysis

Adjusted means for grain yield, grain protein content, and SRI were obtained using residuals derived separately for each environment using the ‘lme4’ function implemented in the R program using the model:

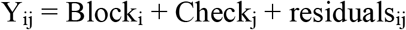

Where Y_ij_ is the phenotypic values of the trait, Check_j_ is the fixed effect of jth replicated check cultivars, and Block_i_ is the fixed effect of the ith block. Residuals were used for obtaining the adjusted means for all the evaluated phenotypic traits (Bates et al., 2015).

Broad sense heritability was extracted using augmented randomized complete block design implemented in R (Aravind et al., 2020) with the model:

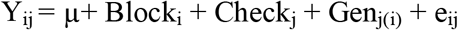

Where Y_ij_, Block_i_, and Check_j_ are defined above, Gen_j(i)_ is the random effect of the unreplicated genotype nested within the ith block and follows the distribution Gen_j_ ~ N(0, σ^2^_g_), and e_ij_ is the standard normal error distributed as e_ij_ ~ N(0, σ^2^_e_).

Genetic correlation between primary traits (grain yield or grain protein content) and secondary traits (individual SRI) are calculated using multivariate models, represented as

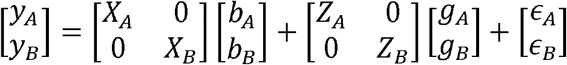

Where *y_A_* and *y_B_* are the BLUPs for the primary (A) and secondary (B) traits, *X* and *Z* denote the design matrix for the fixed and random effect, and *b* is the fixed effects, *g* is the random genetic effects, and *e* is the residuals for each trait. Variance components were calculated assuming 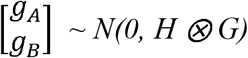, where *G* is the genomic relationship matrix, *H* is the genetic variance-covariance matrix, and 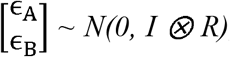, where *R* is the residual variance-covariance matrix, and *I* is the identity matrix. The genetic correlation was obtained as

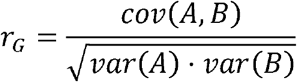

Where *Var(A)*, and *Var(B)* represents the genetic variance of the primary and secondary traits individually and *cov(A, B)* is the covariance between primary and secondary traits. The complete multivariate analysis was performed using a multivariate approach in JMP genomics (SAS Insitute Inc. 2011).

### Genomic prediction models

#### Genomic best linear unbiased predictor (GBLUP)

The UT-GS GBLUP is defined as

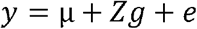

where *y* is the phenotype of interest (grain yield or grain protein content), μ is the overall mean, *Z* is the design matrix linking genotypes to the breeding values, *g* is the vector of genomic breeding values, and *e* is the random residuals. It is assumed that *g* ~ *N(0, Gσ^2^_g_)*, where *G* is genomic relationship matrix, *σ^2^_g_* is the additive genetic variation, and *e* ~ *N(0, Iσ^2^_e_)* with *σ^2^_e_* as residual variance and *I* is the identity matrix. The MT-GS is represented as

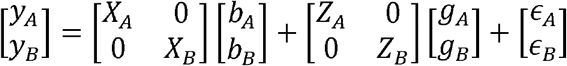

Where *y_A_* and *y_B_* represent the primary and secondary traits, *X* and *Z* are the design matrix associating the fixed and random effects, *b* is the vector of means for primary and secondary traits, *g* and *e* are the vector for random genetic and residual effects. It is assumed as 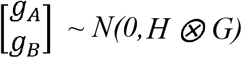, where *G* is the genomic relationship matrix, *H* is the variance-covariance matrix for the two traits, and 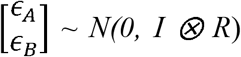, *R* is the residual variance-covariance matrix between two traits (Endelman, 2011; VanRaden, 2008). In all the MT-GS models, individual SRI were included as secondary correlated traits.

#### Bayesian models

As GBLUP uses the relationship matrix for estimating genotypes effects, in this study we also explored the Bayes Lasso, Bayes A, Bayes B, and Bayes Cpi in UT- and MT-GS models, which consider different prior distributions. The UT-GS model is represented as

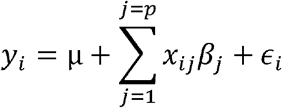

where *y_i_* is the phenotype of interest (grain yield or grain protein content), μ is the overall mean, *β_j_* is the jth marker effect, *x_ij_* is the value of the jth SNP in the ith individual, and *ϵ_i_* is residual error. The conditional prior distribution was separate for each of the Bayesian models employed in this study. For Bayes A, the prior distribution is *ϵ_i_* ~ *N(0, σ^2^)* with *σ^2^* ~ *χ^-2^ (σ^2^/df, S)* for residuals, and *β_j_* ~ *χ^-2^ (df_β_, S_β_)* for genotypic values. Initial values for the degrees of freedom for the t distribution was set to four and *S* was calculated as *S* = *var(y)* * 0.4 as suggested by Pérez & Campos (2014). Analysis was performed using BGLR and MTM packages (Campos & Grüneberg, 2016) with 20,000 Monte Carlo Markov Chain iterations, and 5,000 burn in iterations.

The MTM package was used for fitting MT Bayesian models estimating unstructured variance-covariance between traits. The model is represented as

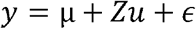

Where *y* is the vector of primary and secondary phenotypic traits, μ is the mean vector for all traits, *u* is the predicted genotypic values for all traits with distribution as *u* ~ *N(0, H ⊗ G)*, where *G* is relationship matrix, *H* is the variance-covariance matrix, and *ϵ* is vector of residuals and distributed as *ϵ* ~ *N(0, I ⊗ R)*, where *I* is the identity matrix and *R* is the variance-covariance matrix for the residuals (Montesinos-López et al., 2016).

#### Random forests (RF)

In RF bootstrap sampling, a subset of features was selected randomly as predictors for splitting the tree nodes. Each tree is chosen for lowering the loss function in the final prediction (Smith et al., 2013). The RF model can be represented as

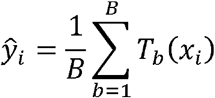

Where *ŷ_i_* is the predicted value of the individual with genotype *x_i_*, *B* is the number of bootstrap samples, and *T* is the total number of trees. RF is computationally less intensive, as each tree is independent of each other and can be computed on different units or nodes (Waldmann, 2016). The working of the RF can be grouped into four main steps:

1. Bootstrap sampling is used to select the individual plant *i (y_i_, x_i_)* with replacement. The sampled individual can appear several times or not, mainly bootstrap sampled *b* = *(1,…, B)*.
2. Selection is performed for the number of features (max features) or input variables at random *(SNP_j_, j = (1,…, J)*, and the best set of features are selected that minimize the loss function obtained as MSE.
3. Splitting is performed at each node into two new subsets (child nodes) for the genotype of *SNP_j_*.
4. Steps 2 and 3 are repeated for each node until a minimum node or the specified max depth is reached. The final predicted value of an individual of genotype *x_i_* is the average of the values predicted by the decision trees in the forest.

The important hyperparameters for RF model training include the number of trees, the number of features sampled for each iteration, the importance of each feature, and the depth of the trees (Hastie et al., 2009). We used randomized and grid search cross-validation for selecting the best hyperparameter’s combination. The combination used for grid search cross-validation after the randomized search was the number of trees (200, 300, 500, 1000), max features (auto, sqrt), and max depth (40, 60, 80, 100). The random forest regressor and Scikit learn libraries were used for analysis in Python 3.7 (Gulli and Pal, 2017).

#### Support vector machine (SVM)

SVM uses kernel functions for mapping the input space into a high dimensional feature space using the relationship between phenotypes and marker genotype (Smola & Scholkopf, 2004). The relationship between the phenotype and genotype is given as

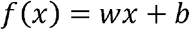

Where *w* is the unknown weight and *b* is the constant, reflecting the maximum allowed bias. The learning of function *f*(*x*) is performed by minimizing the loss function

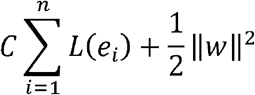

Where *e_i_* = y - *f*(*x*) is the associated error with the *i^th^* training data point, ||*w*||^2^ represents model complexity, *C* is a positive regularization parameter controlling the tradeoff between training error and model complexity, and *L* is the loss function (Vapnik, 2013). Herein, we selected the *ϵ* insensitive loss function for *L* and is represented as

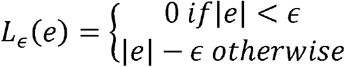

*L* (loss function) is zero if the absolute error is less than predefined *ϵ*, while if absolute error is greater than *ϵ*, *L* is the difference between absolute error and *ϵ*. Usually, *ϵ* insensitive loss function is represented in term of slack variables (*ξξ*\*). The resulting optimization equation can be represented as

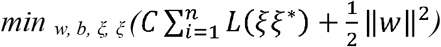

The solution to this minimization problem is of the form

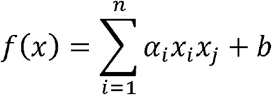

Where *x_i_x_j_* is the inner product of linear function. It was replaced with a non-linear kernel function, namely the gaussian radial basis function. The parameters *C*, *ξ*, and kernel are optimized using cross-validation in the Scikit library (Pedregosa et al., 2011).

#### Multilayer perceptron (MLP)

MLP is a feed-forward neural network consisting of three main layers, namely input, hidden, and output layers. A detailed representation of the MLP used is represented in **Figure 1A**, where the output of a layer depends upon the weighted average transformation of neurons from the previous layer with associated bias. The output of a hidden layer is represented as

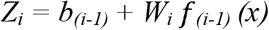

**Figure 1.**
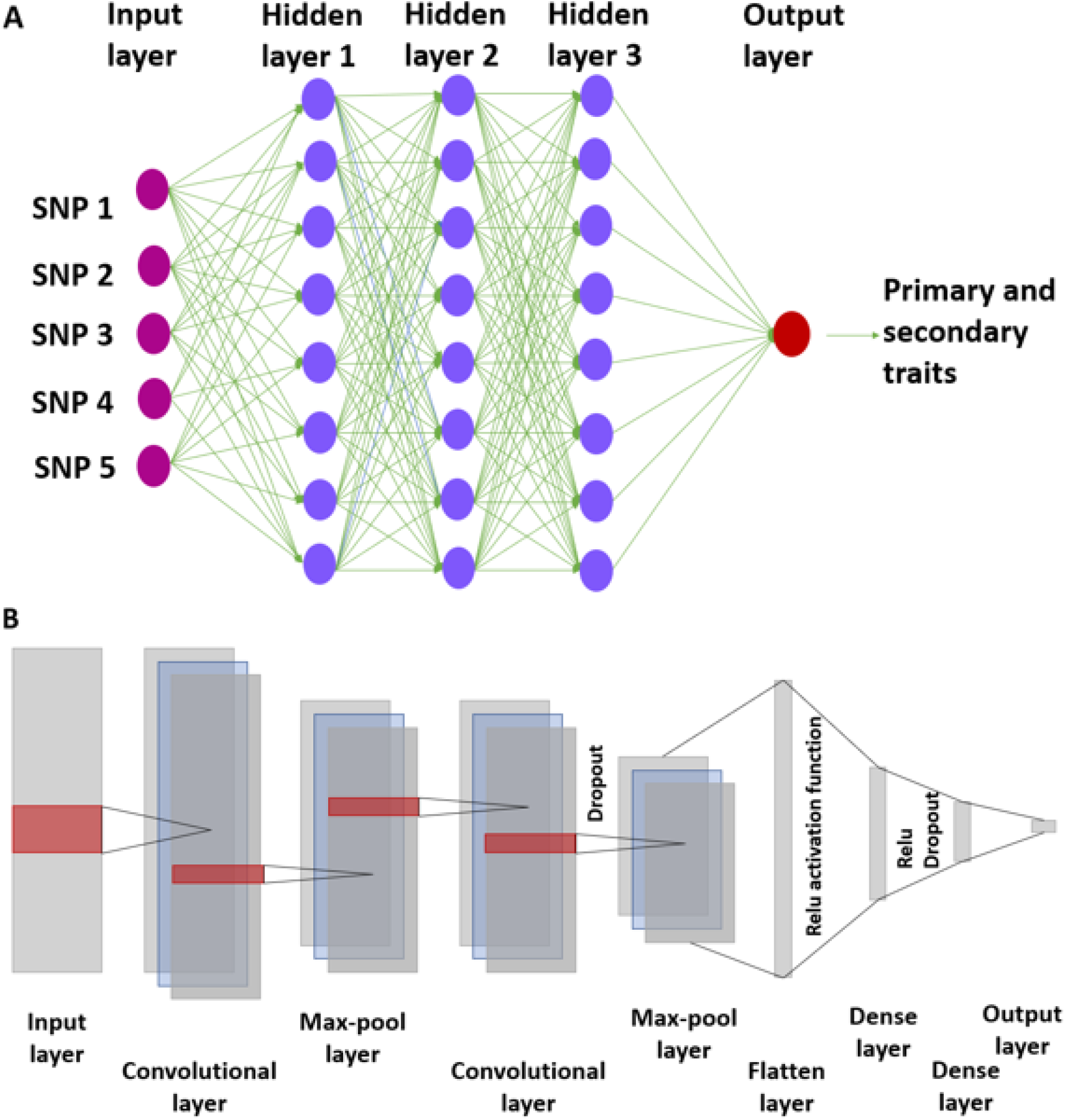
Outline for multilayer perceptron model with one input, three hidden and an output layer. The connections between layers are represented with neurons (A). The representation of convolutional neural network employed in this study is provided with multiple layers (B). Figure 1A was redrawn from code in http://www.texample.net/tikz/examples/neural-network.

Where *Z_i_* is the output from the *i^th^* hidden layer, *W_i_* is the weight associated with the neurons, *f_(i-1)_* represents the activation function linking the associated weights and bias from the previous layer, and this process is repeated until the output layer. In the case of UT-GS models, the output layer is a vector of GEBVs, and in MT-GS, it contains two vectors having GEBVs and spectral information.

Hyperparameters were optimized for the MLP models using inner grid search cross-validation and the Keras function’s internal capabilities. The grid search cross-validation used 80% of the training data, where 80% of this dataset is used for optimizing the hyperparameters and the remaining 20% for validation using Keras independent split validation functions (Cho & Hegde, 2019). The hyperparameters that provided the least MSE on the validation set were selected, and later used on the testing set. Detailed information about the hyperparameter optimization is referred to in a previous publication (Sandhu et al., 2021a). All the MLP algorithms were implemented in Python 3.7 using Keras and Scikit learn libraries (Gulli and Pal, 2017).

#### Convolutional neural network (CNN)

CNN is a special case of neural network used for input features having a specific pattern. The complete layout for CNN is provided in **Figure 1B**. Initial values for the hyperparameters were based on previous findings on UT-GS models (Sandhu et al., 2021a). The CNN model used here consists of one input layer, two convolutional layers, two pooling layers, a dense layer, a flatten layer, two dropouts, and an output layer. Grid search cross-validation was used for selecting hyperparameters, namely, filters (16, 32, 64, 128), learning rate (0.01, 0.05, 0.1), activation function (logistic, linear, tanh, relu), batch size (64, 128), epochs (150, 200), and solver (adam, sgd, lbfgs). These hyperparameters were selected based on previous findings and other studies (Waldmann et al., 2020). Early stopping, dropout, and regularization techniques were applied to control model overfitting. Early stopping involves stopping the training process as validation error reaches a minimum, using Keras-provided API (Callbacks) (Pedregosa et al., 2011). Dropout involves assigning a fixed set of training neurons with a weight to zero for controlling the overfitting and reducing complexity. We used a dropout rate of 0.2 during hyperparameter optimization in MLP and CNN based on Srivastava et al. (2014).

#### Cross-validation and independent prediction

The performances of all nine UT- and MT-GS models were evaluated using five-fold cross-validation. During five-fold cross-validation, 80% of the data was used for model training, and the remaining 20% for model testing within each environment. Two hundred replications were used to assess the model’s performance, and the mean was reported as prediction accuracy. Each replication consisted of five model iterations, where the testing set was rotated for each iteration. Prediction accuracy was obtained as the Pearson correlation coefficient between actual (observed) phenotypic value and the calculated GEBVs. Instant accuracy was reported, which involves the average correlation coefficient of iterations. Comparisons were made between UT- and MT-GS models where a single SRI was included in the MT-GS model. Similarly, machine and deep learning-based MT-GS models were compared with their Bayesian and GBLUP counterparts. MT-GS models used six SRI individually in the model, and the best performing SRI was identified for each trait with different models.

Independent predictions were performed by training models on previous year(s) data and predicting the phenotype in future years. We tested the performance of both UT- and MT-GS with the inclusion of spectral information. In brief, the GS model trained on 2014 and 2015 data was used to predict the 2016 and similarly the model trained on 2014 data to predict the 2015 environment. The GS analysis was computationally intensive, and this problem was resolved by working on Washington State Universities high speed computing cluster (https://hpc.wsu.edu/).

## Results

### Phenotypic summary and heritability

Average phenotypic values and heritabilities are provided for grain yield and grain protein content under three environments (**Table 1**). Grain yield and grain protein content had low and moderate heritability. The six SRI used in this study had moderate to high heritability (**Table 2**), and 2015 had the lowest heritability for phenotypic and spectral traits (**Table 1 and 2**). Phenotypic and genetic correlation between phenotypic traits and SRI was obtained at both heading and grain filling stages (**Table 3, Supplementary Table 2 and 3**). Grain protein content and grain yield had a high and significant correlation with SRI collected at heading and grain filling stage (**Table 3 and Supplementary Table 2**).

**Table 1.**
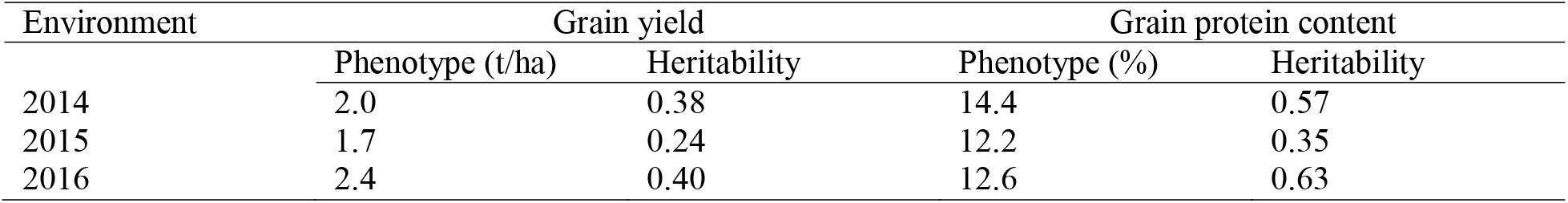
Average phenotypic values and broad sense heritability of grain yield and grain protein content for the three environments (2014-2016).

**Table 2.** Broad sense heritability of six different spectral reflectance indices obtained for each environment for utilization in multi-trait genomic selection models.

| Environment | NDVI<sup>a</sup> | PRI<sup>b</sup> | NWI<sup>c</sup> | ARI<sup>d</sup> | NCPI<sup>e</sup> | GNDVI<sup>f</sup> |
| --- | --- | --- | --- | --- | --- | --- |
| 2014 | 0.75 | 0.93 | 0.60 | 0.74 | 0.64 | 0.69 |
| 2015 | 0.66 | 0.81 | 0.60 | 0.56 | 0.55 | 0.64 |
| 2016 | 0.80 | 0.60 | 0.74 | 0.76 | 0.82 | 0.72 |

<sup>a</sup> NDVI, Normalized difference vegetation index; <sup>b</sup> PRI, Photochemical reflectance index; <sup>c</sup> NWI, Normalized water index; <sup>d</sup> ARI, Anthocyanin reflectance index; <sup>e</sup> NCPI, Normalized chlorophyll pigment ratio index; <sup>f</sup> GNDVI, Green normalized difference vegetation index

**Table 3.**
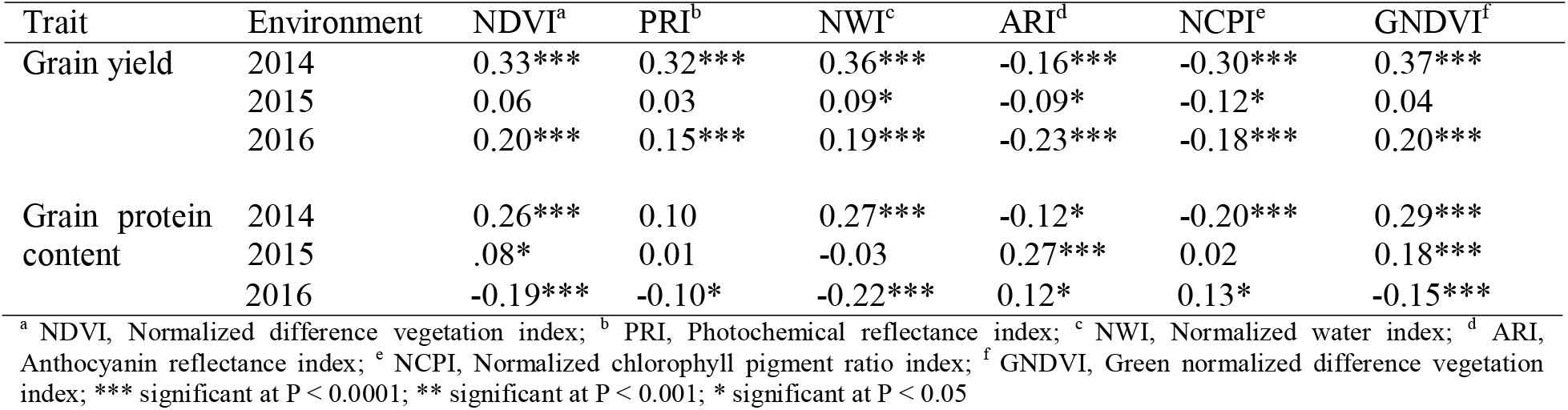
Phenotypic correlation between six different spectral reflectance indices collected at grain filling stage with grain yield and grain protein content for the three environments (2014-2016).

### Genomic selection using uni- and multi-trait models

We evaluated nine different models (GBLUP, Bayes A, Bayes B, Bayes Cpi, Bayes Lasso, RF, SVM, MLP, and CNN) for predicting grain protein content and grain yield using five-fold cross-validation (**Figure 2 and 3**). A single SRI was included in each model for predicting both traits under MT-GS models and their average results are depicted for comparison with UT-GS models (**Figure 2 and 3**). In the case of grain yield, MT-GS models either gave an equal or higher prediction accuracy than UT-GS models (**Supplementary Table 4**). Furthermore, machine and deep learning models performed better than the traditional GBLUP and Bayesian models under UT and MT models. The improvement in prediction accuracy with MT-GS models for grain yield varied from 0 to 28.5%, with maximum improvement observed in the 2014 environment, and the lowest increase was observed for the 2015 environment (**Figure 2**). RF and MLP performed best for predicting grain yield under all the environments, closely followed by GBLUP (**Supplementary Table 4 and Figure 2**). Both models performed superior for UT- and MT-GS models compared to other machine and deep learning models. Four bayesian models, namely, Bayes A, Bayes B, Bayes Cpi, and Bayes Lasso, produced almost the same prediction accuracy for grain yield under the UT- and MT-GS models (**Supplementary Table 4 and Figure 2**). SVM resulted in the lowest prediction accuracy for grain yield and CNN observed the lowest increase under the MT-GS models (**Figure 2 and Supplementary Table 4**).

**Figure 2.**
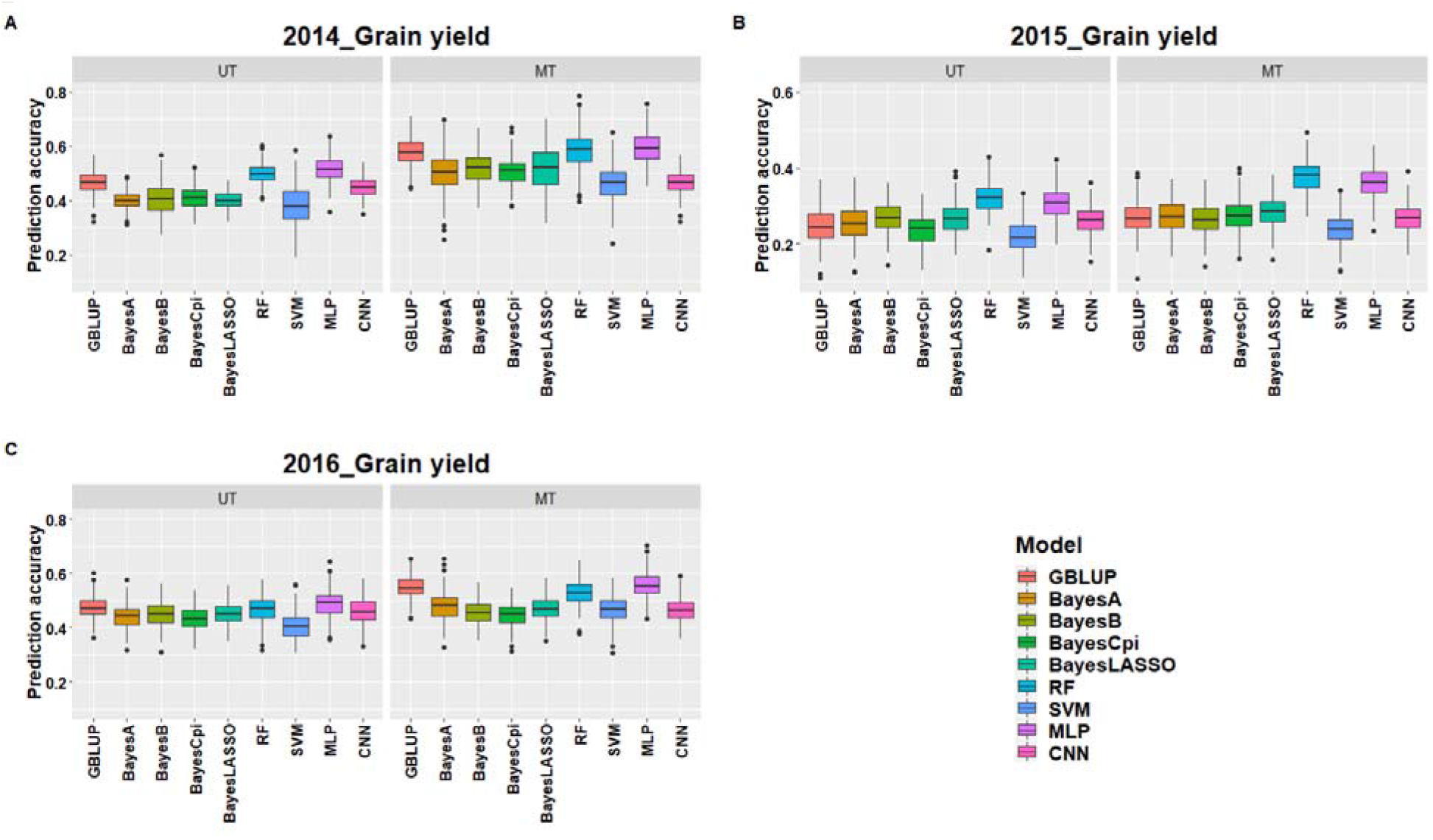
Prediction accuracies for grain yield with nine different uni- and multi-trait genomic selection models under the three different environments (2014-16) (**A-C**) using five-fold cross-validation. The x-axis represents the nine genomic selection models with faceting separating the uni- and multi-trait models.

**Figure 3.**
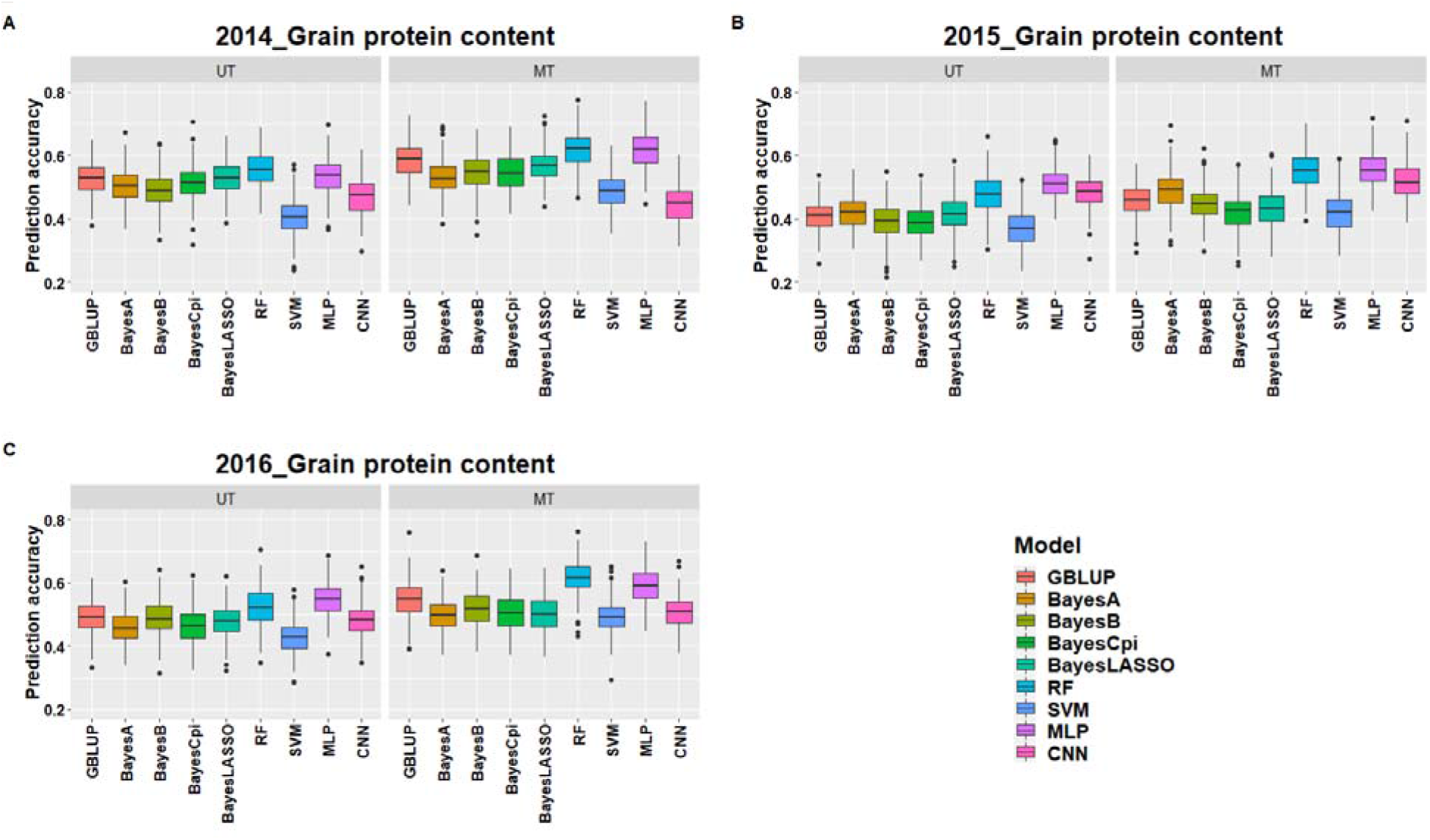
Prediction accuracies for grain protein content with nine different uni- and multi-trait genomic selection model under the three different environments (2014-16) (**A-C**) using five-fold cross-validation. The x-axis represents the nine genomic selection models with faceting separates the uni- and multi-trait models.

Similarly, MT-GS models increased prediction accuracy for grain protein content compared to UT-GS counterparts (**Figure 3 and Supplementary Table 4**). MLP and RF performed superior to other models and gave similar accuracy under different scenarios (**Figure 3 and Supplementary Table 4**). The performance of MT-GS models varied from −0.04 to 15% compared to the UT-GS models. Similar to grain yield, SVM performed poorest for UT- and MT-GS models to predict grain protein content. We observed similar trends in improvement in prediction accuracy with MT-GS models for all environments compared to grain yield, with 2014 observing the largest increase (**Figure 2 and 3**). The maximum prediction accuracy for grain protein content was 0.62 in 2014 with RF, 0.56 in 2015 with MLP, and 0.61 in 2016 with RF MT-GS models (**Figure 3**). We observed that similar to grain yield, MT-GS model for CNN resulted in the lowest increase in prediction accuracy.

### Performances of multi-trait genomic selection models using individual SRI

We evaluated each SRI’s relationship with grain protein content and grain yield across all nine MT-GS models assessed in this study (**Figure 4 and 5**). Inclusion of any SRI in the MT-GS model resulted in higher prediction accuracy than the UT-GS model for grain protein content and grain yield. **Figure 4** shows the prediction accuracies for grain protein content for the three environments with six individual SRIs in the MT-GS models. GNDVI was the best performing index for seven out of the nine models in each environment (**Figure 4**). RF and MLP resulted in the greatest improvement in prediction accuracy by including GNDVI in the MT-GS model. SVM was the only model where GNDVI performed worse than other indices for most environments (**Figure 4**). The maximum prediction accuracy for grain protein content in 2014, 2015, and 2016 was 0.67, 0.61, and 0.68, respectively, with RF by inclusion of GNDVI in the MT-GS model.

**Figure 4.**
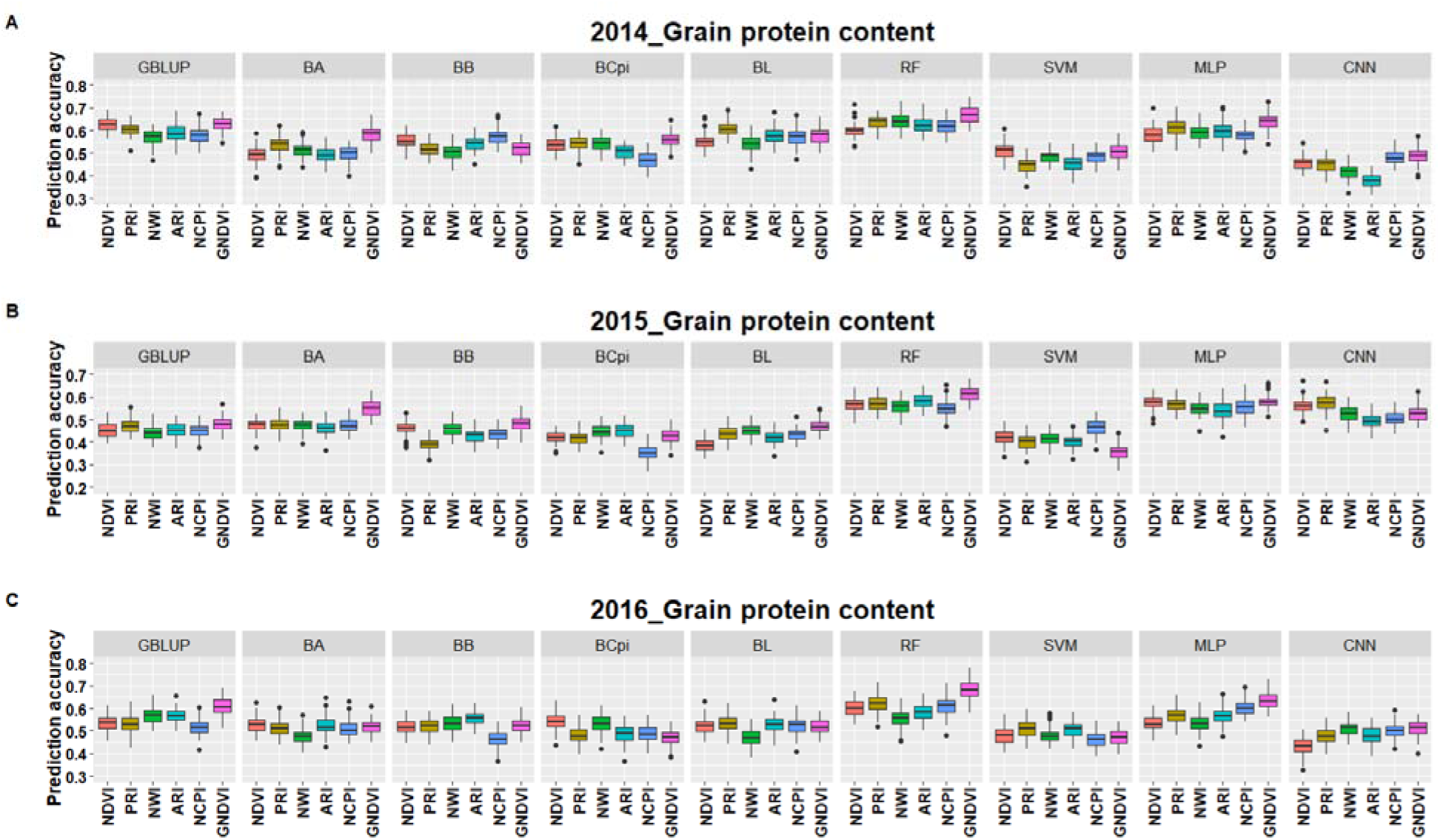
Prediction accuracies for grain protein content for the three environments (**A-C**) with inclusion of six different spectral reflectance indices in the nine multi-trait genomic selection models. The x-axis represents the individual spectral reflectance indices and multi-trait genomic selection models are separated with facets for comparing across model performances.

**Figure 5.**
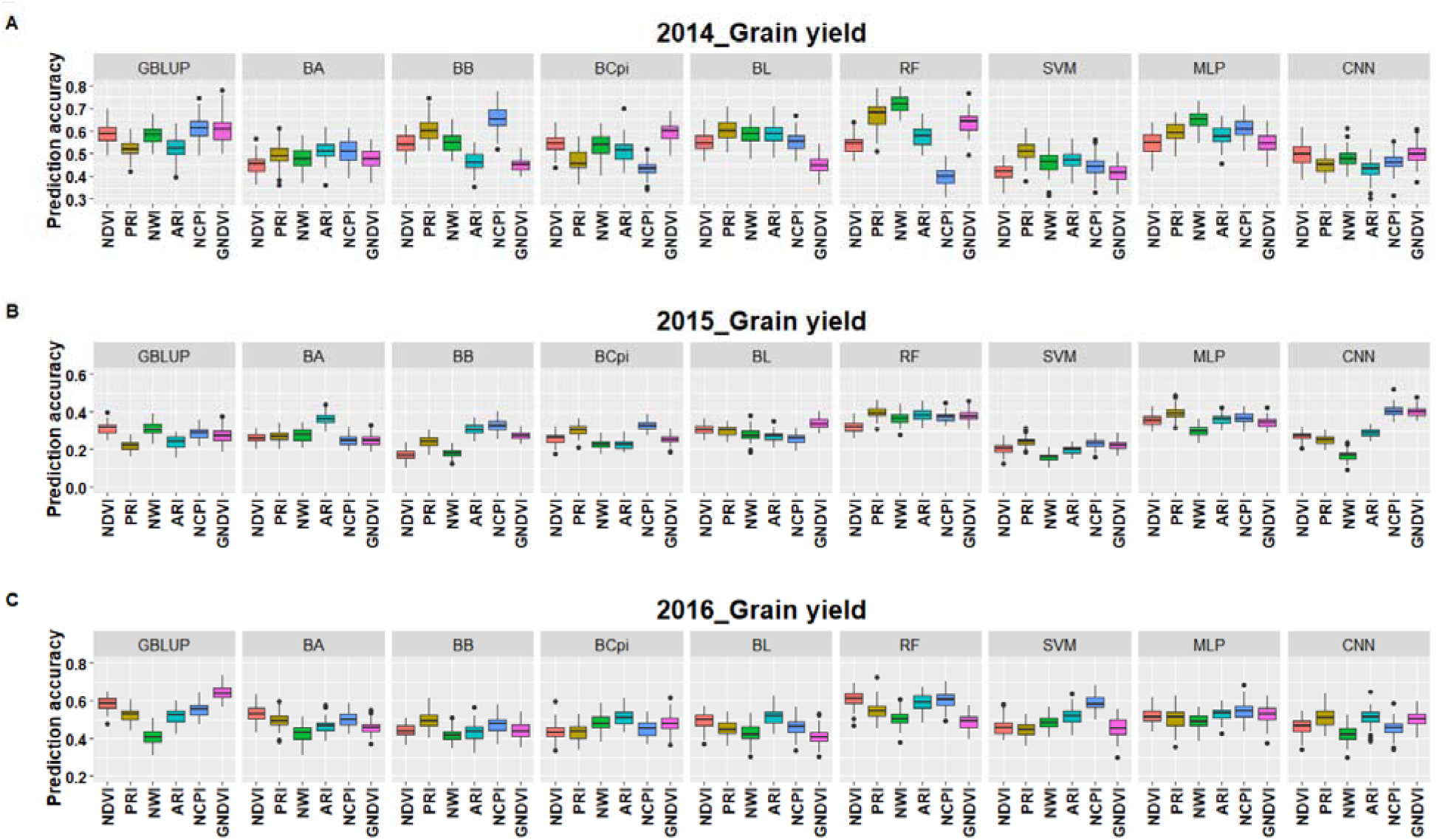
Prediction accuracies for grain yield for the three environments (**A-C**) with inclusion of six different spectral reflectance indices in the nine multi-trait genomic selection models. The x-axis represents the individual spectral reflectance indices and multi-trait genomic selection models are separated with facets for comparing across model performances.

Similarly, the inclusion of individual SRI in the MT-GS models increased prediction accuracy compared to UT-GS models for grain yield (**Figure 5**). Unlike grain protein content, there was no individual SRI that performed best across all environments and models. The maximum prediction accuracy for 2014 was 0.72 with inclusion of NWI, 2015 was 0.40 with inclusion of NDVI, and 2016 was 0.64, with inclusion of NDVI. Overall, we observed that NWI, NDVI, and GNDVI provide the necessary information for increasing prediction accuracy for grain yield (**Figure 5**).

### Independent predictions with uni- and multi-trait genomic selection models

Independent predictions involve prediction across environments where models were trained on previous year datasets, and predictions were made for upcoming years. We used four UT- and MT-GS models, namely, GBLUP, RF, MLP, and CNN, for the independent predictions as they performed consistently better with cross-validation for both traits and under all the environments. SVM was excluded because of its poor performance for both traits. All the Bayesian models performed similarly, but had less accuracy, so were excluded from independent predictions because of huge computational time limitations. The machine and deep learning models were approximately four times faster than the Bayesian models. **Figure 6** shows the results for independent prediction for both traits using UT- and MT-GS models.

**Figure 6.**
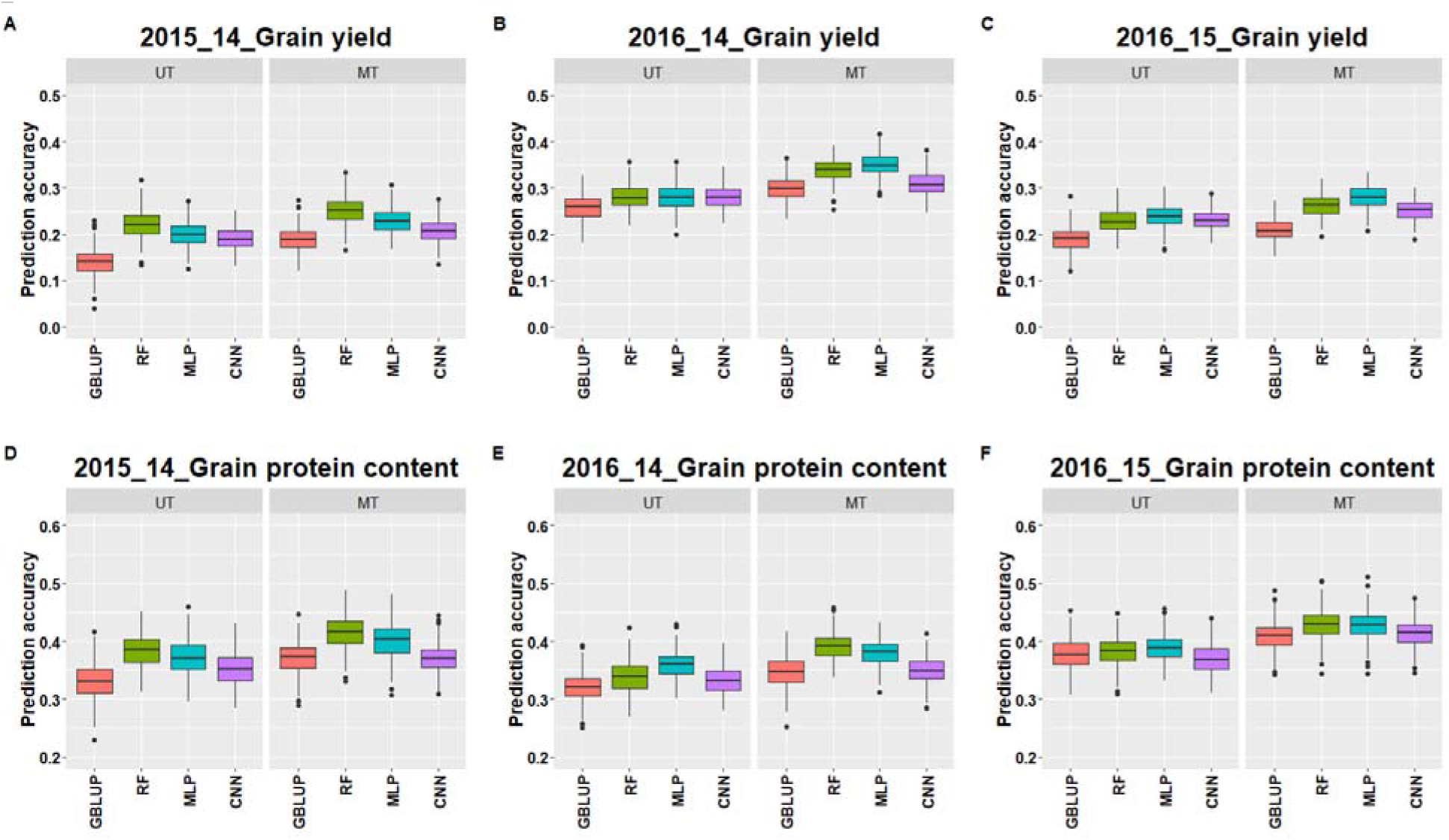
Independent prediction accuracies for grain yield (**A-C**) and grain protein content (**D-F**) using four different uni- and multi-trait genomic selection models. The first digit of the year represents the testing environment, and the second year represents the training environment. The x axis represents the different models and faceting separate the uni- and multi-trait models.

There was an increase of 17% and 11% in prediction accuracies for grain yield and grain protein content with MT-GS over the UT-GS models (**Supplementary Table 5**). RF and MLP performed consistently better under UT- and MT-GS models to predict grain yield and grain protein content, closely followed by CNN. The highest average prediction for grain yield and grain protein content was 0.29 and 0.41 with MLP and RF, compared to 0.20 and 0.34 with the traditional UT GBLUP model (**Supplementary Table 5**). There was a varied increment in prediction accuracies with different MT-GS models for grain yield and grain protein content, as depicted in **Figure 6**.

## Discussion

Grain yield and grain protein content are highly important target traits in wheat breeding programs and for other cereal grains. The generally negative correlation between them, along with lower heritability, creates a problem in efficiently selecting both the traits simultaneously. GS and high throughput phenotyping have the potential for reducing the challenges associated with selection for these two traits. GS may increase selection efficiency, reduce the generation advancement time, and increase selection intensity (Heffner et al., 2010). Similarly, spectral information from phenomics data allows for indirect selection by using SRI as a proxy indicator. The increased use of machine and deep learning models in other disciplines has prompted its use in plant breeding. Several MT selection studies have conducted and demonstrated the potential for increasing prediction accuracy (Bhatta et al., 2020; Jia & Jannink, 2012). This study evaluated the potential of MT machine and deep learning models for predicting grain yield and grain protein content in wheat using spectral information as a secondary trait. The spectral information acts as a proxy indicator for selection, is correlated with the primary trait of interest, and has higher heritability. We observed that machine and deep learning models, namely RF and MLP, resulted in an increased prediction accuracy for both traits when spectral information was included in MT-GS models.

### Trait characterization and association

The greatest advantage of MT-GS models is borrowing the information from secondary traits to increase prediction accuracies for the primary trait (Calus & Veerkamp, 2011). Understanding the genetic architecture of each traits is the first consideration when developing MT models. The primary traits of interest in this study were grain yield and grain protein content, which were evaluated for three years and heritabilities were low to moderate (**Table 1**), suggesting a considerable influence of non-genetic effects. Lower heritability for grain yield and grain protein content was expected, as these traits are controlled by a large number of small effect QTLs and are genetically complex. Similar results were obtained in previous studies for grain yield and grain protein content (Sun et al., 2017). Secondary traits (SRI used in this study) have high heritability and correlate positively with the primary traits (**Table 2**). Furthermore, the collection of SRI is easy and could be performed using high-throughput techniques (Sankaran et al., 2015). This suggested that their inclusion in MT-GS models may improve prediction accuracy, increase selection intensity, and lead to faster breeding cycles (Crain et al., 2018).

GNDVI was the best performing index for seven out of nine models evaluated in this study, and could be a useful proxy index for selecting grain protein content in breeding programs. Association of GNDVI with nitrogen status and translocation is related to the increase in prediction accuracies for grain protein content in the MT-GS models (Gitelson et al., 1996). The high correlation of primary traits with SRI indicates a direct connection between them. Grain protein content has a lower heritability value than GNDVI, and hence a lesser amount of variation is accounted by GS models for grain protein content under the UT-GS model. The higher accuracy observed in MT-GS models can be attributed to capturing more genetic variation, as GNDVI is genetically correlated with grain protein content. Grain protein content had the highest correlation with GNDVI, which measures reflection from the green region (550 nm) of the plant vegetation spectrum and provides information about the plant’s nitrogen status (Gitelson et al., 1996). Another advantage of using GNDVI is that it allows the prediction of grain protein content earlier in the selection pipeline, saving the time, cost, and effort to harvest and collect data from a large number of field plots. Using GNDVI also aids in improving MT-GS models, which can be used early in the selection pipeline to select improved lines for advancement in the breeding program.

Grain yield is a complex trait resulting from a myriad of interactions, including nutrient status, water availability, biotic and abiotic stress. Three SRI, namely NWI, NDVI, and GNDVI, resulted in the highest prediction accuracy for grain yield under the MT-GS models. These indices each measure a part of NIR (900-970 nm), and this spectrum determines the water status and biomass of the plants, suggesting NIR is useful for predicting grain yield, especially in the Pacific Northwest US, where wheat is grown under dryland conditions. Identifying multiple SRI that increase prediction accuracy provided insight that model analysis should not rely on a single SRI in each year. Expression of several physiological processes in plants is dependent upon the plants genetic makeup, factors like light, temperature, humidity, day length, etc, and the growth stage when the plant experiences stress. Different SRI are able to capture these various genotype by environment interactions, along with environmental variation that may exist from year to year, which contribute to final grain yield estimations. The inclusion of multiple SRI in the model helps explain the unknown variance component ignored in UT-GS models. We were able to identify the three SRI that are more influential for predicting grain yield out of the six SRI explored. Using these three SRI will reduce computation time and cost for data management to make selections.

### Potential of the machine and deep learning models in a breeding program

This study aimed to explore machine and deep learning potential in wheat breeding programs using MT-GS models. Deep learning is a new machine learning branch using a dense network of neurons to explore the dataset’s complicated hidden relationship. We concluded that MT random forest and multilayer perceptron resulted in an improvement of 23-31% prediction accuracy for both traits under cross-validation and independent prediction compared to the GBLUP and Bayesian models. Similar results were obtained by Ma et al. (2018) for predicting grain yield, plant height, and grain length in wheat using UT-deep learning models and rrBLUP. Their study demonstrated potential for the utilization of deep learning models in plant breeding. Deep learning based GS models gave 0-5% higher prediction accuracies for various agronomic traits in wheat in our previous work (Sandhu et al., 2021a). Montesinos-López et al. (2018b) concluded that deep learning models were superior for six out of nine traits evaluated in maize and wheat over the traditional GBLUP. Additionally, Montesinos-López et al. (2018a) showed that MT deep learning models performed superior to the MT Bayesian models when genotype by environment interaction was not included for predicting grain yield in wheat. These results open up a new path for improved breeding selection, that could translate into higher rates of genetic gain.

Machine and deep learning models, unlike Bayesian models, are highly flexible for mapping complex interactions present between predictors and responses, and thereby interpreting the trend of the current dataset (Liu et al., 2019). Bayesian models often include selection of a specific subset of markers which explain the most variation in the response, in contrast to machine and deep learning models, that explore the whole data space during model training (Pérez & Campos, 2014). As grain yield and grain protein content are polygenic, relying on Bayesian models might not be a good strategy, as it is often unknown how many QTLs are present in a particular population and are expressing under the specific environment. Hence, machine and deep learning models are suitable for complex traits as they explore all possible relationships between markers and traits. Machine and deep learning models also account for interactions among predictors and remove redundant information using filters, nodes, or neurons (Crossa et al., 2019), and modelling of this interaction is especially important for MT models when primary and secondary traits are correlated. Our results confirm this, obtaining the highest prediction accuracy with a MT-MLP model, compared to a UT-MLP model. The neuron weight updates in the hidden layers leads to a combination of attributes for capturing the most suitable hierarchical representation. Similarly, random forest development uses independent tree branches, and the conclusion is based on the forest’s average with individual tree branches being uncorrelated, instead of any one individual branch (Waldmann, 2016). In this way, both random forest and MLP models are an improvement for mapping complex interactions and resulted in the highest prediction accuracy for both grain protein content and grain yield in this study.

The computation time increased linearly with inclusion of more variables such as SRI in this study for MT GBLUP and Bayesian models; however, MT machine and deep learning models are well developed for parallelism in computation (Lecun et al., 2015). We observed that MT machine and deep learning models were four times faster than the MT Bayesian models, due to their capacity to parallelize the operations. With the continuous increase in secondary traits owing to utilization of high throughput phenotyping tools in the breeding program, breeders need to shift from traditional Bayesian models to more computationally efficient models, and machine and deep learning models provide a new avenue in this regard. Grain yield and grain protein content had different heritability in each environment, with 2015 being lowest. The MT machine and deep learning models resulted in the maximum increase in prediction accuracy for both traits in the 2015 environment, showing that these models are also superior for lower heritability traits or environments, which is often seen in plants (González-Camacho et al., 2018).

It is a misconception that machine and deep learning models should be used only on large datasets having thousands of individuals, which is difficult for traits like grain yield or grain protein content which are evaluated at mid to late stages in breeding programs (Angermueller et al., 2016). However, our results and other related work indicate that machine and deep learning models have similar or higher performance compared to traditional GBLUP and Bayesian models when using data from hundreds of individuals for model training (Zingaretti et al., 2020). Furthermore, a large dataset with 100k individuals was used for predictions with deep learning models in the GS context and did not observe any superiority over the traditional linear mixed models (Bellot et al., 2018). Zingaretti et al. (2020) and Montesinos-López et al. (2018a) demonstrated deep learning models have higher prediction accuracy than GBLUP and other mixed models when using 1233 strawberry and 250 wheat genotypes. These results, combined with ours, suggest that training population size plays a minor role compared to the genetic architecture of the trait and secondary traits utilized in the models. However, adequate training population sizes remain important in GS.

The main issue with machine and deep learning models is the lack of biological significance as different hyperparameters in the models handle different data parts. These models might not be useful in providing genetic insight for the primary and secondary traits employed, and hence genome-wide association studies are an important complement. Furthermore, compared to GBLUP, machine and deep learning models were computationally intensive because of the extra step of hyperparameter optimization. Hyperparameter optimization is required separately for each trait and could significantly discourage users when results are needed quickly in breeding programs for making selections (Cho & Hegde, 2019). Plant breeders are often interested in each predictor’s significance in the models, which is not possible in deep learning because of their black-box nature due to many hidden layers, neurons, and filters. Finally, machine and deep learning model implementation requires a sufficient background in computer science, mathematics, and machine learning, which will require additional efforts by plant breeders, or accomplished through efficient collaborations. Although there are some potential hindrances in implementing machine and deep learning models, their utilization can result in the improvement of prediction accuracy for complex traits of interest in breeding programs. Overall, this study presents the benefits to utilizing MT machine and deep learning models for predicting complex traits under selection while efficiently using spectral information.

### Conclusion

In this study, we evaluated the potential of MT machine and deep learning models for predicting grain yield and grain protein content using spectral information in a wheat breeding program. The model’s performances were compared with traditional UT and MT GBLUP and Bayesian models under cross-validation and independent predictions. Our results showed the vast potential for MT machine and deep learning models with spectral information as a proxy phenotype. Random forest and multilayer perceptron were the best performing models for both traits under all the evaluated scenarios, closely followed by convolutional neural network and GBLUP. Green normalized difference vegetation index was the best SRI for predicting grain protein content for most MT models under cross-validation and independent predictions. Furthermore, machine and deep learning models were competitive with Bayesian models as they were less computationally intensive than Bayesian models. This and previous studies on deep learning for predicting complex traits shows enormous potential for the utilization of these models in plant breeding programs to enhance genetic gain for quantitative traits.

## Supporting information

Not applicable

## Abbreviations

ARI: anthocyanin reflectance index
CNN: convolutional neural network
GBLUP: genomic best linear unbiased predictor
GEBVs: genomic estimated breeding values
GNDVI: green normalized difference vegetation index
GS: genomic selection
MLP: multilayer perceptron
MT: multi-trait
NCPI: normalized chlorophyll pigment ratio index
NDVI: normalized difference vegetation index
NWI: normalized water index
PRI: photochemical reflectance index
RF: random forest
SVM: support vector machine
UT: uni-trait

## Authors contributions

KS: conceptualized the idea, analyzed data, and drafted the manuscript; SP: assisted in data analysis and edited the manuscript; MP: conducted field trials, edited the manuscript and obtained the funding for the project; AC: supervised the study, conducted field trials, edited the manuscript and obtained the funding for the project.

