## Supplementary material for "Multi-Trait Machine and Deep Learning Models for Genomic Selection using Spectral Information in a Wheat Breeding Program": Not applicable

| **Supplementary Table 1.** Information about the six spectral reflectance indices (SRI) used in this study. | | | |
| --- | --- | --- | --- |
| Index | Formula^+^ | Physiological processes | References |
| Normalized difference Vegetation Index (NDVI) | (R^800 _^R^680^)/  (R^800^+R^680^) | Biomass, plant health, vegetative greenness | (Rouse et al., 1972) |
| Photochemical reflectance index (PRI) | (R^531_^R^570^)/  (R^531^+R^570^) | Carotenoid content | (Peñuelas et al., 1997) |
| Normalized water index (NWI) | (R^970^-R^880^)/  (R^970^+R^880^) | Plant water status | (Prasad et al., 2007) |
| Anthocyanin reflectance index (ARI) | R^800^  (1/R^550_^1/R^700^) | Anthocyanin pigment | (Gitelson et al., 1996) |
| Normalized chlorophyll pigment ratio index (NCPI) | (R^680_^R^430^)/  (R^680^+R^430^) | Chlorophyll pigments | (Peñuelas et al., 1994) |
| Green normalized difference vegetation index (GNDVI) | (R^780_^R^550^)/  (R^780^+R^550^) | Chlorophyll content | (Gitelson et al., 1996) |
| + R represents the reflection at a particular wavelength | | | |

| **Supplementary Table 2.** Phenotypic correlation between six different spectral reflectance indices collected at heading stage with grain yield and grain protein content for the three environments (2014-2016). | | | | | | | |
| --- | --- | --- | --- | --- | --- | --- | --- |
| Trait | Environment | NDVI^a^ | PRI^b^ | NWI^c^ | ARI^d^ | NCPI^e^ | GNDVI^f^ |
| Grain yield | 2014 | 0.20*** | 0.21*** | 0.19*** | -0.16*** | -0.13* | 0.26*** |
|  | 2015 | 0.03 | 0.07 | 0.00 | -0.09* | -0.07 | 0.05 |
|  | 2016 | 0.15*** | 0.16*** | 0.05 | -0.23*** | -0.15*** | 0.20*** |
| Grain protein content | 2014 | 0.35*** | 0.19*** | 0.35*** | -0.12* | -0.38*** | 0.34*** |
|  | 2015 | 0.19*** | 0.11* | 0.21*** | 0.27*** | -0.19*** | 0.28*** |
|  | 2016 | -0.20*** | 0.02 | -0.16*** | 0.12* | 0.08 | -0.20*** |
| ^a^ NDVI, Normalized difference vegetation index; ^b^ PRI, Photochemical reflectance index; ^c^ NWI, Normalized water index; ^d^ ARI, Anthocyanin reflectance index; ^e^ NCPI, Normalized chlorophyll pigment ratio index; ^f^ GNDVI, Green normalized difference vegetation index; *** significant at P < 0.0001; ** significant at P < 0.001; * significant at P < 0.05 | | | | | | | |

| **Supplementary Table 3.** Genetic correlation of six different spectral reflectance indices with grain yield and grain protein content. | | | | | | |
| --- | --- | --- | --- | --- | --- | --- |
| Trait | NDVI^a^ | PRI^b^ | NWI^c^ | ARI^d^ | NCPI^e^ | GNDVI^f^ |
| Grain yield | 0.73 | 0.52 | 0.65 | 0.59 | 0.56 | 0.65 |
| Grain protein content | 0.61 | 0.48 | 0.65 | 0.55 | 0.53 | 0.70 |
| ^a^ NDVI, Normalized difference vegetation index; ^b^ PRI, Photochemical reflectance index; ^c^ NWI, Normalized water index; ^d^ ARI, Anthocyanin reflectance index; ^e^ NCPI, Normalized chlorophyll pigment ratio index; ^f^ GNDVI, Green normalized difference vegetation index; all genetic correlations are significant at p < 0.05 | | | | | | |

| **Supplementary Table 4.** Average prediction accuracies for grain yield and grain protein content under the cross-validation for three environments (2014-2016) by the nine univariate-trait (UT) and multi-trait (MT) GS models. The highest prediction accuracy for each scenario is bolded. | | | | | | | | | | |
| --- | --- | --- | --- | --- | --- | --- | --- | --- | --- | --- |
| Trait |  | GBLUP | BayesA | BayesB | BayesCpi | BL | RF | SVM | MLP | CNN |
| Grain yield | UT | 0.38 | 0.36 | 0.38 | 0.36 | 0.37 | 0.43 | 0.33 | **0.44** | 0.39 |
|  | MT | 0.47 | 0.41 | 0.42 | 0.41 | 0.43 | **0.50** | 0.39 | 0.49 | 0.40 |
| Grain protein content | UT | 0.47 | 0.46 | 0.46 | 0.46 | 0.48 | 0.52 | 0.40 | **0.53** | 0.48 |
|  | MT | 0.53 | 0.51 | 0.51 | 0.49 | 0.50 | **0.59** | 0.47 | **0.59** | 0.49 |

| **Supplementary Table 5.** Average prediction accuracies for grain yield and grain protein content under three different scenarios of independent predictions using four different univariate-trait (UT) and multi-trait (MT) GS models. The highest prediction accuracy for each case is bolded. | | | | | |
| --- | --- | --- | --- | --- | --- |
| Trait |  | GBLUP | RF | MLP | CNN |
| Grain yield | UT | 0.20 | **0.24** | **0.24** | 0.23 |
|  | MT | 0.23 | 0.28 | **0.29** | 0.26 |
| Grain protein content | UT | 0.34 | **0.37** | **0.37** | 0.35 |
|  | MT | 0.38 | 0**.41** | 0.40 | 0.38 |
